# Occurrence cubes: a new paradigm for aggregating species occurrence data

**DOI:** 10.1101/2020.03.23.983601

**Authors:** Damiano Oldoni, Quentin Groom, Tim Adriaens, Amy J.S. Davis, Lien Reyserhove, Diederik Strubbe, Sonia Vanderhoeven, Peter Desmet

## Abstract

In this paper we describe a method of aggregating species occurrence data into what we coined “occurrence cubes”. The aggregated data can be perceived as a cube with three dimensions - taxonomic, temporal and geographic - and takes into account the spatial uncertainty of each occurrence. The aggregation level of each of the three dimensions can be adapted to the scope. Built on Open Science principles, the method is easily automated and reproducible, and can be used for species trend indicators, maps and distribution models. We are using the method to aggregate species occurrence data for Europe per taxon, year and 1km^2^ European reference grid, to feed indicators and risk mapping/modelling for the Tracking Invasive Alien Species (TrIAS) project.

## Introduction

To address the ongoing biodiversity crisis policymakers demand rapid, reliable and regular information on the status of biodiversity. The Group on Earth Observations Biodiversity Observation Network (GeoBON) have proposed a suite of Essential Biodiversity Variables (EBV) that are intended to be a minimal set of variables required to report biodiversity change (Pereira et al., 2013). To create such EBVs for species distribution and abundance it has been proposed to create aggregated “data cubes” with taxonomic, spatial and temporal dimensions (Kissling et al., 2018). The concept is that such cubes can be generated automatically and repeatedly from raw observation data as often as needed. Nevertheless, although pilot projects have created workflows to EBVs on a small scale, no one has actually shown how such a cube can be generated in a manner that does not require considerable manual intervention (Hardisty et al., 2018, 2019).

In the recent decades the volume of published occurrence data has increased enormously, partly thanks to research infrastructures such as the Global Biodiversity Information Facility (GBIF) (www.gbif.org), but also the digitization of legacy data and the use of mobile applications for recording (Blagoderov et al., 2012; Chandler et al., 2017; Pocock et al., 2019). These occurrences are extensively used in ecology for many purposes such as species distribution modelling, risk mapping and calculating biodiversity indicators of extent and spread. However, an unavoidable problem with the use of such massive volumes of data is their heterogeneity. Occurrences have an intrinsic spatial uncertainty, which is not always negligible. This heterogeneity is the result of the numerous methods used to collect data and the wide range of people doing the observing. They can be collected in many ways, such as ecological surveys of single sites, gridded data created for national atlases, or casual observations from citizen scientists.

Typical techniques to deal with spatial uncertainty is either removing insufficiently precise occurrences or gridding occurrences in cells that encompass the highest expected uncertainty, effectively not using the spatial uncertainty associated with each occurrence. The method we describe in this paper does take into account this spatial uncertainty.

Occurrences can be defined as objects in a three-dimensional space where the dimensions are:

1. Taxonomic: what
2. Temporal: when
3. Spatial: where

Occurrences can be aggregated along all three dimensions.

The taxonomic dimension is categorical, so in principle aggregation is optional. However, the presence of synonyms makes the aggregation process relevant. Research infrastructures such as GBIF and OBIS use a taxonomic backbone so that occurrences of synonyms are automatically associated with the corresponding accepted taxon, thus making aggregation at species level or higher ranks relatively easy. The same holds true for occurrences of infraspecific taxa: they are automatically returned when searching for occurrences of species or higher rank.

The temporal dimension is a continuum. Several Darwin Core standard (DwC) terms are available to provide standardized temporal information. The most important field is `eventDate` (http://rs.tdwg.org/dwc/terms/eventDate), defined as the “date-time or interval during which an Event occurred”. Occurrences are generally easy to aggregate as temporal uncertainty is typically considerably lower than the aggregation span used in the vast majority of the applications, typically year.

The spatial dimension is also continuous and aggregation is theoretically possible by using reference grids, such as the European reference grid of the European Environment Agency (EEA) or the United States National Grid of the Federal Geographic Data Committee (FGDC). However, spatial constraints are needed as all reference grids are local to avoid distortions due to projecting the curved Earth on a flat surface. Moreover, conceiving occurrences as points with infinite precision is not only unrealistic but also incorrect and misleading. Some atlas datasets, for example, are already aggregated using a different grid and/or scale, which means that all occurrences are assigned to the centroids of the grid cells. Neglecting this aspect could result in an underestimation of the area of occupancy as severe as the overestimation of the abundance in the cell grids the centroids belong to. In this paper we state that an occurrence is spatially representable as a closed plane figure such as a circle or a polygon, never as the geometric center (centroid) of it. The most used way to express spatial uncertainty is by using a radius which defines, together with geographical coordinates (e.g. latitude/longitude), a circle, although atlas data could be better described as squares (Figure 1).

**Figure 1.**
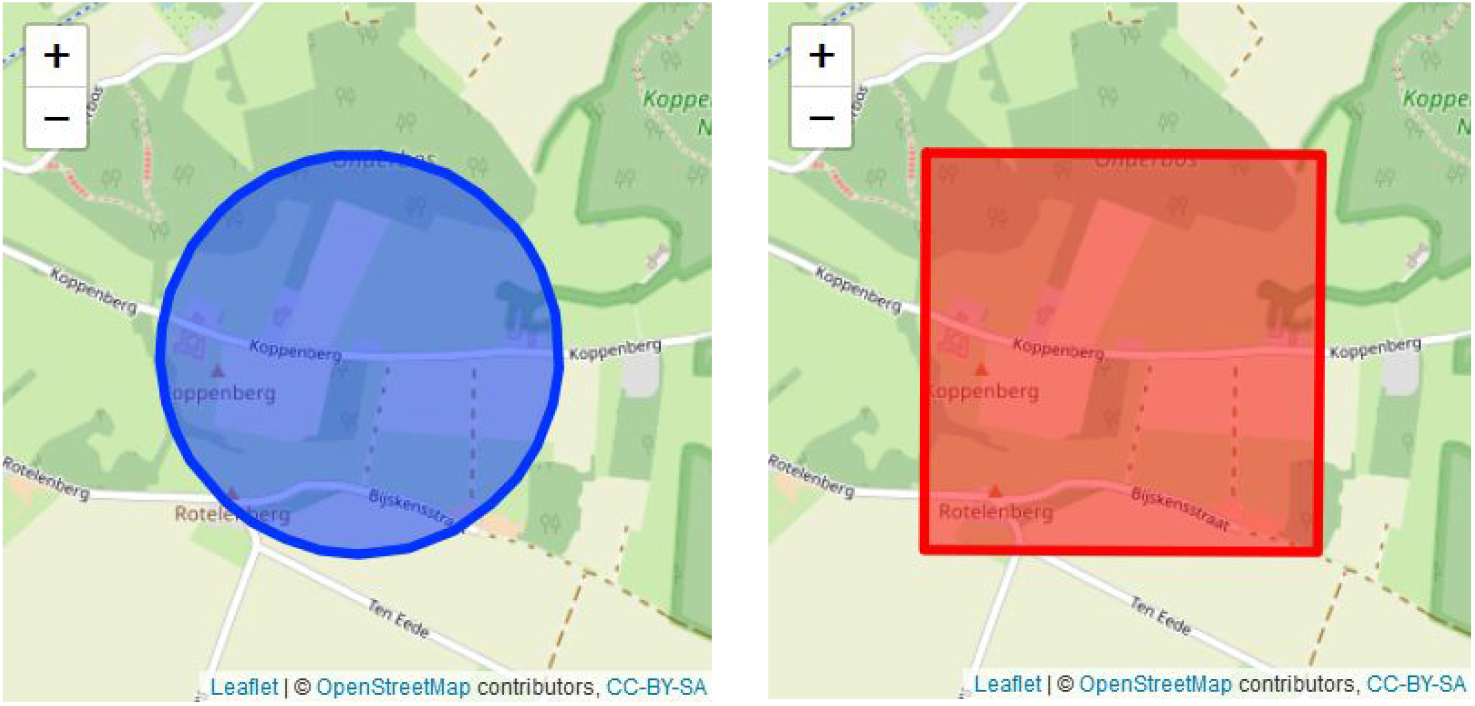
Left: occurrences from GPS are circles. The stronger the GPS signal, the smaller the circle. Right: occurrences from atlas data are squares.

Darwin Core Standard (DwC) provides different terms to express spatial uncertainty of an occurrence in a consistent way: `coordinatePrecision` (http://rs.tdwg.org/dwc/terms/coordinatePrecision), `coordinateUncertaintyInMeters` (http://rs.tdwg.org/dwc/terms/coordinateUncertaintyInMeters) and `pointRadiusSpatialFit` (http://rs.tdwg.org/dwc/terms/pointRadiusSpatialFit). The `coordinatePrecision` defines the decimal precision of the coordinates given in the fields `decimalLatitude` and `decimalLongitude`. The `coordinateUncertaintyInMeters` defines the radius describing the smallest circle containing the whole of the location. This way of expressing uncertainty became popular with the advent of GPS receivers as these instruments define a location as a circle, with a radius depending on the sensibility of the signal. Uncertainty of occurrences in atlas data are squares. For this reason Darwin Core standard has a `pointRadiusSpatialFit` term, defined as “The ratio of the area of the point-radius (decimalLatitude, decimalLongitude, coordinateUncertaintyInMeters) to the area of the true (original, or most specific) spatial representation of the Location.”

In case of occurrences collected at a scale of *lxl m*^2^ the `coordinateUncertaintyInMeters` is typically defined as the radius of the circumscribed circle with radius 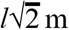, while the `pointRadiusSpatialFit` is the ratio of this circle’s area, *l*^2^ · π/2 *m*^2^, to the grid square area, *l*^2^ *m*^2^, i.e. π/2. As for GBIF occurrence data, the `coordinateUncertaintyInMeters` is the most used term for storing spatial uncertainty, even for gridded data. In this paper we will explain how the spatial uncertainty expressed by the `coordinateUncertaintyInMeters` can be used to produce aggregated occurrence data. For the remainder of this document we will refer to occurrences as circles, even if the method described belows general and can be applied to any plane figure used to represent the occurrences.

## Methodology

The production of what we call “occurrence cubes” can be divided in four steps:

1. Specify taxonomic, temporal and geographical constraints and granularity
2. Harvest occurrences and quality assessment
3. Assign occurrences spatially by taking into account their coordinate uncertainty
4. Aggregate occurrences along taxonomic, temporal and spatial dimension into an occurrence cube

We illustrate the methodology below with a minimal, reproducible example: an occurrence cube for GBIF occurrences of the genus *Reynoutria* in Belgium from 2000 to 2018, aggregated by species, year and 1km EEA reference grid cell

### Step 1. Specify constraints and granularity

We have first to delimitate the “space” along the taxonomic, temporal and spatial dimensions, by applying some constraints. Such constraints are typically defined by the scope of the research. Are you interested in the occurrences of taxa within kingdom Animalia or within class Aves? Are you interested in occurrences related to a specific time window? Is your research limited to a specific country or a specific area? Example: genus *Reynoutria* from 2000 to 2018 in Belgium.

However, defining constraints is not enough. Aggregation requires a discretization of the delimited space. The level of granularity resulting from such discretization depends on the scope of the research as well. Example: by species, year and reference grid at 1km scale provided by the European Environment Agency (EEA).

The **taxonomic** dimension is by definition not continuous and occurrences can be identified at different taxonomic ranks. The granularity of the taxonomic dimension is therefore the rank at which we want to aggregate. The taxonomic backbone of research infrastructures such as GBIF and Ocean Biogeographic Information System (OBIS) automatically returns all the occurrences of taxa with lower ranks and link them to the higher ranks. The same holds true for synonyms as well: a taxonomic backbone helps to solve synonymy as all the occurrences of synonyms point to the corresponding accepted taxon. In our example, we want to aggregate by species. For e.g. *Reynoutria japonica*, GBIF will automatically include occurrences associated with that accepted name, as well as two synonyms and the infraspecific variety *Fallopia japonica var. japonica* These four scientific names all share *Reynoutria japonica* in the `species` field (as shown in Table 1) making aggregation easier.

**Table 1.** Taxon of occurrences of species *Reynoutria japonica* as returned by GBIF. Occurrences of synonyms and infraspecific taxa are returned as well and all share the accepted species in the field `species`.

| scientificName | taxonRank | species | taxonomicStatus |
| --- | --- | --- | --- |
| Reynoutria japonica<br>Houtt. | SPECIES | Reynoutria japonica | ACCEPTED |
| Fallopia japonica (Houtt.)<br>Ronse Decraene | SPECIES | Reynoutria japonica | SYNONYM |
| Fallopia compacta<br>(Hook.fil.) G.H.Loos &<br>P.Keil | SPECIES | Reynoutria japonica | SYNONYM |
| Fallopia japonica var.<br>japonica | VARIETY | Reynoutria japonica | DOUBTFUL |

The **time** of an occurrence is typically defined with a precision way higher than the granularity used for aggregation. For being statistically significant, EBVs are built using a temporal resolution of at least one year.

As regards the **spatial** granularity, we encourage, where possible, the use of standard reference grids provided and maintained by governmental institutions. For Europe we can use the EEA reference grid system, a set of grids at 1km, 10km and 100km scale for each European country. Aside from the scope of the research, the spatial granularity depends at a certain extent on the area defined by the spatial constraint. As shown in Figure 2, the occurrence cube of Luxemburg at the scale of 100km is not so useful while the same cannot be said about the occurrence cube of a vast country as France.

**Figure 2.**
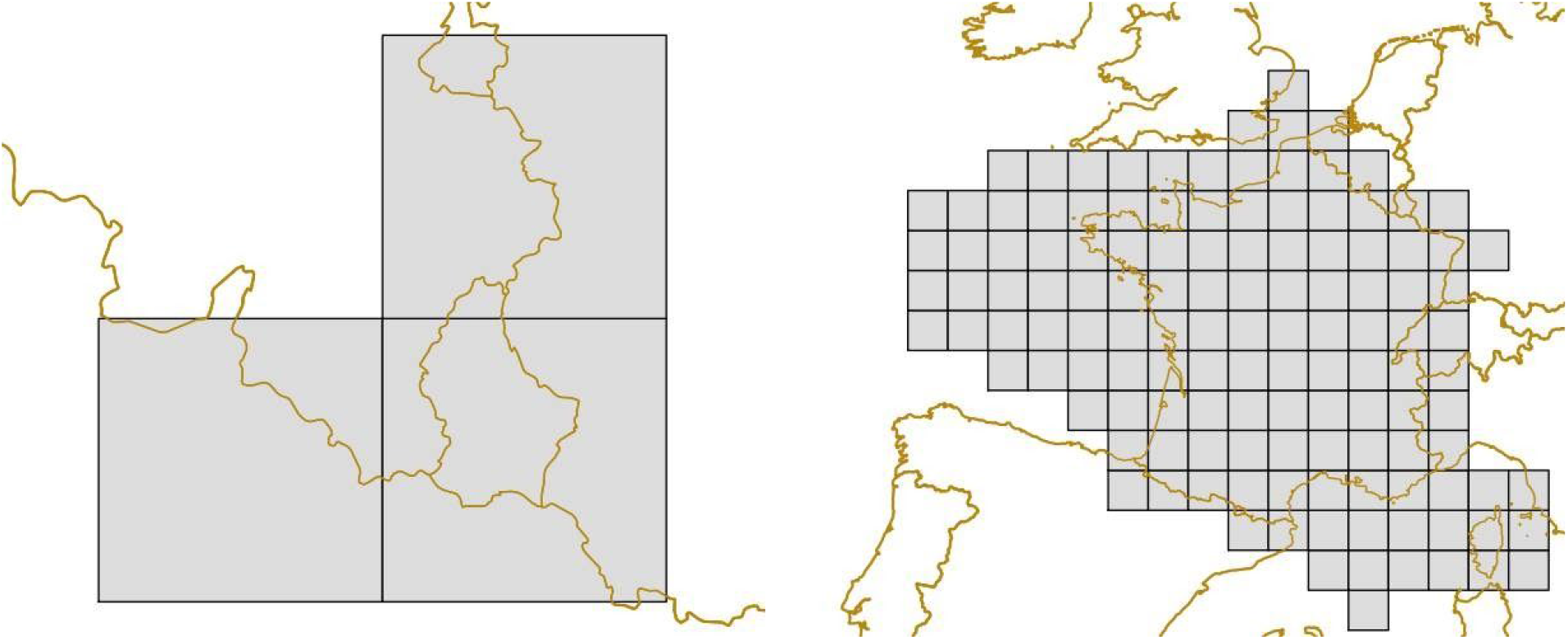
European Environment Agency (EEA) reference grids at 100km scale of Luxemburg (left) and France (right). Source: EEA.

### Step 2. Harvest occurrences and quality assessment

The sources for harvesting occurrences are typically world-wide research infrastructures, like GBIF and OBIS. Data from such infrastructures are open and standardized. The more sources you choose, the more preprocessing is needed, especially if some data sources are not standardized. To make FAIR occurrence cubes, it is also important to make harvested occurrence data findable. For example, in case of GBIF data this is possible as a triggered download gets an unique Digital Object Identifier (DOI). Example: the DOI of our download is https://doi.org/10.15468/dl.aobecp and contains 36,851 occurrences.

Some basic data quality assessment is needed to remove invalid occurrences, e.g. occurrences with invalid or suspicious coordinates, occurrences related to fossil or living specimens or occurrences representing absences. Depending on the infrastructure of the data source, it is possible to add the data-quality checks defined above directly in the download query, thus reducing data volume. Additional data screening can be applied, e.g. removing data coming from a particular dataset considered not reliable enough or data with suspicious event dates.

However, occurrences without spatial *uncertainty* should be screened. Often the spatial uncertainty can be inferred from the metadata, although it can be impractical as the occurrences can be harvested from many different datasets. One could discard these data, but then potentially loses useful and quite precise casual observations as well. We opt for assigning a default spatial uncertainty of 1000 meter to these occurrences, thus creating circles with a radius of 1000 m. In our example, the field `coordinateUncertaintyInMeters` is missing for 450 occurrences. These occurrences come from two citizen science projects and a sampling event dataset of a public research institution. Not discarding them, but assigning instead a default spatial uncertainty of 1000 m seems the most reasonable option. The result of this process is shown in Figure 3.

**Figure 3.**
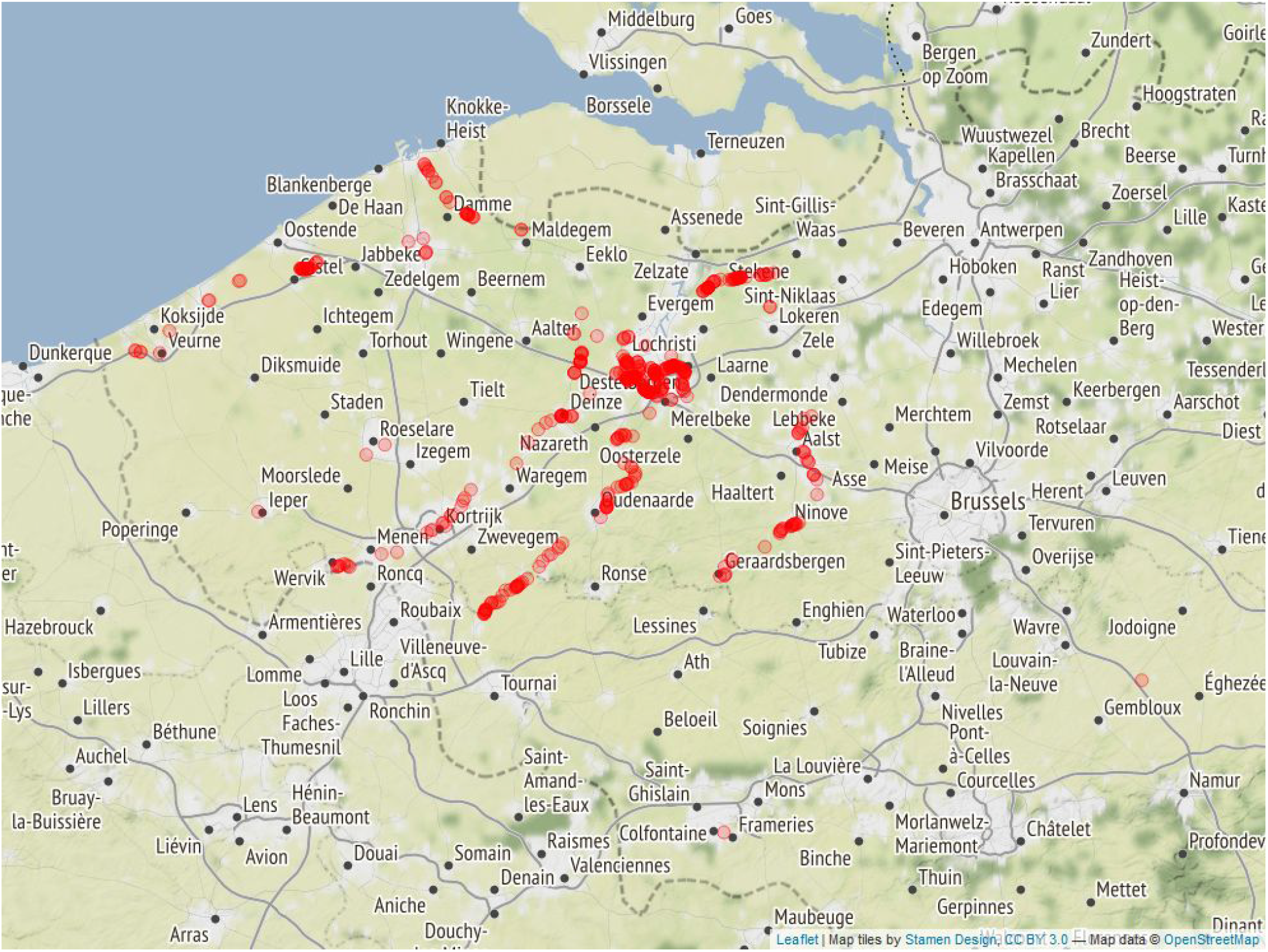
Spatial uncertainty is set to 1000 meters for the 450 occurrences of genus *Reynoutria* where the value of field `coordinateUncertaintyInMeters` is missing.

### Step 3. Assign occurrences to a reference grid

Once all occurrences have a valid spatial uncertainty, we can now assign them to the cells of the reference grid. Geometrically this operation sounds like: how to assign a circle to squares? The answer is trivial only if the circle representing the occurrence is completely contained in one cell. Since that is often not the case, we propose to randomly choose a point within the circle and assign the occurrence to the cell this point belongs to. In Figure 4 we show some cases found in our example.

**Figure 4.**
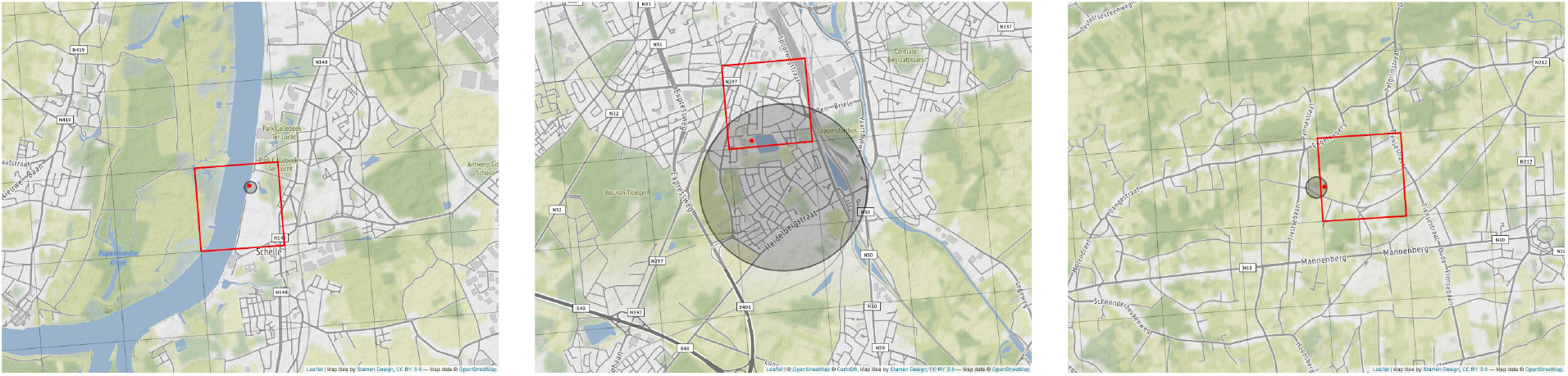
A random point (red point) is chosen within the circle (gray) defining the occurrence. The occurrence is then assigned to the cell the point belongs to (red square). Left: an occurrence (https://www.gbif.org/occurrence/2235280677) is totally contained in one of the cells of the reference grid. Center: An occurrence (https://www.gbif.org/occurrence/1569856810) spreads over multiple cells. Right: an occurrence (https://www.gbif.org/occurrence/2235279067) with small uncertainty spreads over two cells.

The probability distribution is by definition uniform all over the circle so the probability that the random point falls in a specific cell is equal to the proportion of the circle’s area covered by the cell. Geometrically it can be demonstrated that no cell has a higher probability to get the occurrence assigned than that one containing the center. However, this doesn’t exclude the possibility that the occurrence would be assigned to another cell as shown in Figure. 4.

### Step 4. Aggregate occurrences

Aggregating occurrences means *counting* how many occurrences of a specific taxon are in a specific cell and in a specific time interval. Using our example with occurrences of *Reynoutria*, where we decided to produce an occurrence cube at species and year level using a reference grid at 1km resolution, we have to count how many occurrences of *Reynoutria* are there within each year, cell and species. As the occurrence cubes can be used as input for modelling and risk assessment, we store the smallest geographic coordinate uncertainty of the occurrences assigned to a certain cell as value as well. Using a tabular structure (typical of R *data.frames* or pandas *DataFrames*), an occurrence cube would look like a table with as many columns as the sum of the number of dimensions (three) and the number of values (two). In Table 2 we show an excerpt from the example occurrence cube.

**Table 2.** Tabular representation of the occurrence cube of *Reynoutria* in Belgium from 2000 to 2018. The first three columns represent the temporal, spatial and taxonomic dimensions respectively. Column `year` contains the year the occurrences took place, `eea_cell_code` the cell code from the EEA reference grid at 1km scale, `speciesKey` the GBIF identifier of the species (2889173: *Reynoutria japonica*, 4038485: *Reynoutria bohemica*, 2889088: *Reynoutria sachalinensis*). Taxonomic-spatial-temporal triplets with no occurrences are omitted.

| year | eea_cell_code | speciesKey | n | min_coord_uncertainty |
| --- | --- | --- | --- | --- |
| 2000 | 1kmE3809N3113 | 2889173 | 1 | 700 |
| 2000 | 1kmE3809N3135 | 2889173 | 1 | 700 |
| ... | ... | ... | ... | ... |
| 2006 | 1kmE3936N3071 | 2889173 | 1 | 49 |
| 2006 | 1kmE3947N3132 | 2889088 | 1 | 700 |
| ... | ... | ... | ... | ... |
| 2010 | 1kmE3883N3121 | 4038485 | 1 | 700 |
| 2010 | 1kmE3884N3121 | 2889173 | 1 | 10 |
| ... | ... | ... | ... | ... |
| 2014 | 1kmE3886N3121 | 2889173 | 51 | 10 |
| 2014 | 1kmE3886N3122 | 2889173 | 109 | 10 |
| ... | ... | ... | ... | ... |
| 2018 | 1kmE4047N3067 | 2889173 | 1 | 2828 |

As mentioned in Step 1, defining the taxonomic granularity of the occurrence cube implies that occurrences linked to a taxon can come from multiple taxa such as synonyms or taxa with lower rank. For this reason, it can be informative to provide a taxonomic compendium of the occurrence cube as shown in Table. 3. The full occurrence cube and the taxonomic compendium are available on GitHub: https://github.com/trias-project/occurrence-cube-paper/tree/9426a29dc6f080920509aa295bd49dad0ea10d26/data/processed.

**Table 3.** Taxonomic compendium of the occurrence cube from GBIF occurrences of genus *Reynoutria*, in Belgium from 2000 to 2018. As shown in column `includes`, occurrences of a species can come from synonyms or infraspecific taxa, described by their GBIF taxon keys and scientific names.

| speciesKey | species | includes |
| --- | --- | --- |
| 2889088 | *Reynoutria sachalinensis* | 5334293: *Fallopia sachalinensis* (Friedrich Schmidt Petrop.) Ronse Decraene<br>2889088: *Reynoutria sachalinensis* Nakai |
| 2889173 | *Reynoutria japonica* | 5334357: *Fallopia japonica* (Houtt.) Ronse Decraene<br>2889173: *Reynoutria japonica* Houtt.<br>8361333: *Fallopia compacta* (Hook.fil.) G.H.Loos & P.Keil<br>7128523: *Fallopia japonica* var. *japonica* |
| 4038485 | *Reynoutria bohemica* | 5652296: *Fallopia bohemica* (Chrték & Chrtková) J.P.Bailey<br>4038485: *Reynoutria bohemica* Chrték & Chrtková |

## The occurrence cube

The resulting occurrence cube can be projected on an orthogonal plane by aggregating along one of the three dimensions, as shown in Figure 5a.

**Figure 5.**
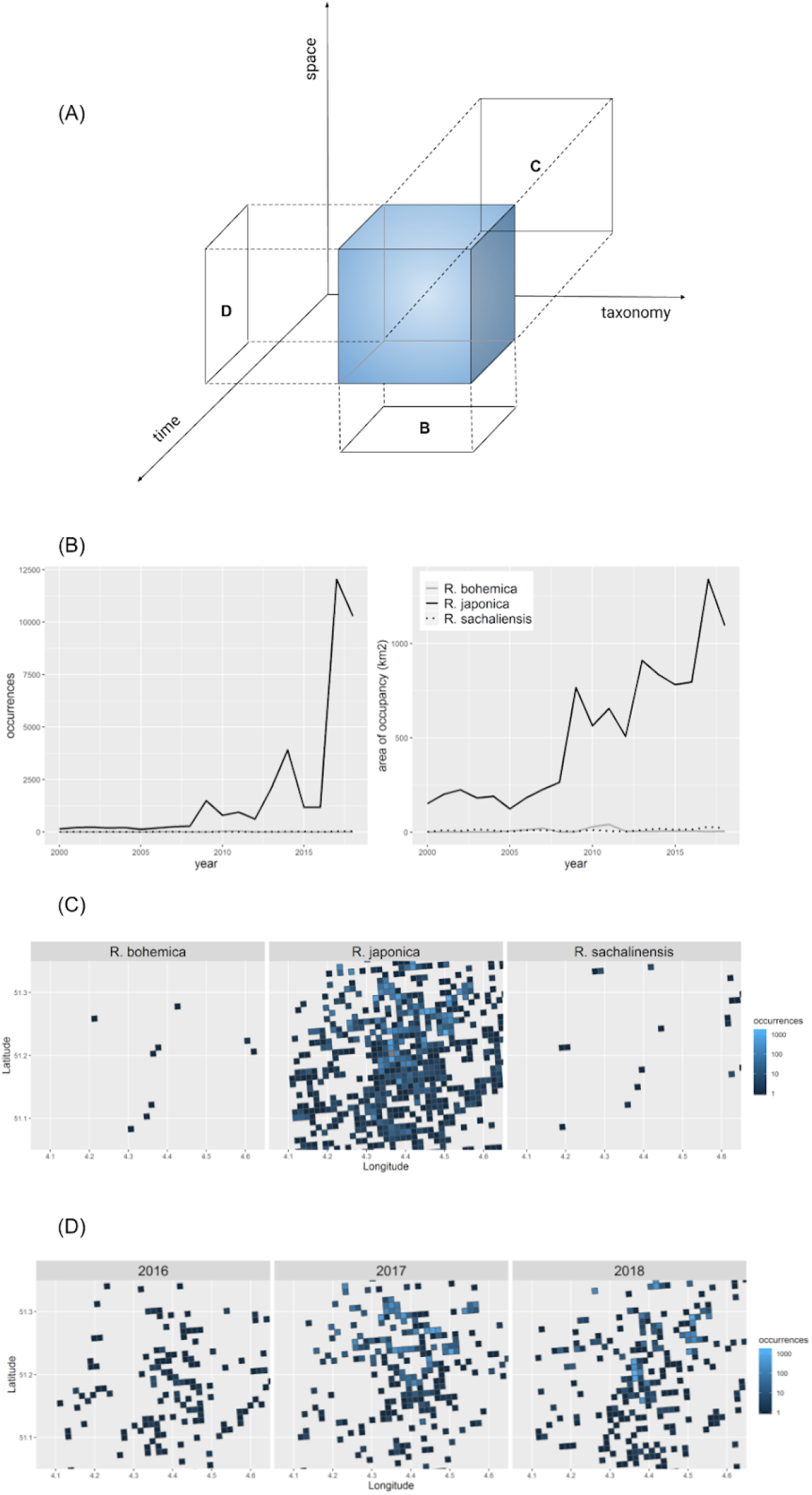
(A) The occurrence cube and its projections on the temporal/taxonomic plane (B), the taxonomic/spatial plane (C) and the temporal/spatial plane (D). (B) Number of occurrences (left) and number of 1×1km cells or *area of occupancy* (right) of *Reynoutria bohemica, R. japonica* and *R. sachalinensis* per year. Both indicators can be seen as ways of projecting the occurrence cube on the temporal/taxonomic dimensions. (C) Projecting the occurrence cube along the taxonomic/spatial plane, thus getting a heatmap of the number of occurrences for each of the *Reynoutria sp*. in Belgium. The maps are zoomed for better readability. (D) Projecting the occurrence cube along the temporal/spatial plane, thus getting a heatmap of the number of occurrences of genus *Reynoutria* in Belgium for each year. The maps are zoomed for better readability.

Aggregating along the spatial dimensions means projecting the cube on the taxonomic and temporal dimensions. Counting the number of occurrences we get the abundance, counting the number of occupied cells we get the area of occupancy as shown in Figure 5b. Aggregating along the temporal dimension means projecting the cube on the spatial and taxonomic dimensions. Based on our example, it means counting how many occurrences of each species of *Reynoutria* are in each cell during the entire period 2000 - 2018, as shown in Figure 5c. Similarly, aggregating along the taxonomic dimension, we project the cube on the spatial and temporal dimensions. We are then counting the number of occurrences of genus *Reynoutria* per cell and year as shown in Figure 5d.

We applied this methodology to larger taxonomic, spatial and temporal constraints as well. We created and published occurrence cubes at species level for Belgium and Italy (Oldoni et al., 2020a) and the occurrence cubes for non-native taxa in Belgium and Europe (Oldoni et al., 2020b). All these occurrence cubes are at year level and are based on EEA reference grids at 1km scale.

## Notes

Data, scripts and figures are open and available on GitHub: https://github.com/trias-project/occurrence-cube-paper

## Acknowledgements

This work has been funded under the Belgian Science Policies Brain program (BelSPO BR/165/A1/TrIAS).

## Notes

https://github.com/trias-project/occurrence-cube-paper

https://zenodo.org/record/3637911

https://zenodo.org/record/3635510

## References

Blagoderov, V., Kitching, I.J., Livermore, L., Simonsen, T.J., & Smith, V.S. (2012). No specimen left behind: industrial scale digitization of natural history collections. ZooKeys, (209), 133. https://doi.org/10.3897/zookeys.209.3178

Chandler, M., See, L., Copas, K., Bonde, A.M., López, B.C., Danielsen, F., Legind, J.K., Masinde, S., Miller-Rushing, A.J., Newman, G., Rosemartin, A., & Turak, E. (2017). Contribution of citizen science towards international biodiversity monitoring. Biological Conservation, 213, 280–294. https://doi.org/10.1016/j.biocon.2016.09.004

Hardisty, A.R., Belbin, L., Hobern, D., McGeoch, M.A., Pirzl, R., Williams, K.J., & Kissling, W.D.D. (2018). Research infrastructure challenges in preparing essential biodiversity variables data products for alien invasive species. Environmental Research Letters. https://doi.org/10.1088/1748-9326/aaf5db

Hardisty, A.R., Michener, W.K., Agosti, D., García, E.A., Bastin, L., Belbin, L., Bowser, A., Buttigieg, P.L., Canhos, D.A.L., Egloff, W., De Giovanni, R., Figueira, R., Groom, Q., Guralnick, R.P., Hobern, D., Hugo, W., Koureas, D., Ji, L., Los W., Manuel, J., Manset, D., Poelen, J., Saarenmaa, H., Schigel, D., Uhlir, P.F., & Kissling, W.D. (2019). The Bari Manifesto: An interoperability framework for essential biodiversity variables. Ecological informatics, 49, 22–31. https://doi.org/10.1016/j.ecoinf.2018.11.003

Kissling, W.D., Ahumada, J.A., Bowser, A., Fernandez, M., Fernández, N., García, E.A., Guralnick, R.P., Isaac, N.J.B., Kelling, S., Los, W., McRae, L., Mihoub, J.-B., Obst, M., Santamaria, M., Skidmore, A.K., Williams, K.J., Agosti, D., Amariles, D., Arvanitidis, C., Bastin, L., De Leo, F., Egloff, W., Elith, J., Hobern, D., Martin, D., Pereira, H.M., Pesole, G., Peterseil, J., Saarenmaa, H., Schigel, D., Schmeller, D.S., Segata, N., Turak, E., Uhlir, P.F., Wee, B., & Hardisty, A.R. (2018), Building essential biodiversity variables (EBVs) of species distribution and abundance at a global scale. Biol Rev, 93: 600–625. https://doi.org/10.1111/brv.12359

Oldoni, D., Groom, Q., Adriaens, T., Davis, A.J.S., Reyserhove, L., Strubbe, D., Vanderhoeven, S., & Desmet, P. (2020a). Occurrence cubes at species level for European countries (Version 20200205) [Data set]. Zenodo. http://doi.org/10.5281/zenodo.3637911

Oldoni, D., Groom, Q., Adriaens, T., Davis, A.J.S., Reyserhove, L., Strubbe, D., Vanderhoeven, S., & Desmet, P. (2020b). Occurrence cubes for non-native taxa in Belgium and Europe (Version 20200204) [Data set]. Zenodo. http://doi.org/10.5281/zenodo.3635510

Pereira, H.M., Ferrier, S., Walters, M., Geller, G.N., Jongman, R.H.G., Scholes, R.J., Bruford, M.W., Brummitt, N., Butchart, S.H.M., Cardoso, A.C., Coops, N.C., Dulloo, E., Faith, D.P., Freyhof, J., Gregory, R.D., Heip, C., Höft, R., Hurtt, G., Jetz, W., Karp, D.S., McGeoch, M.A., Obura, D., Onoda, Y., Pettorelli, N., Reyers, B., Sayre, R., Scharlemann, J.P.W., Stuart, S.N., Turak, E., Walpole, M., & Wegmann, M. (2013). Essential biodiversity variables. Science, 339(6117), 277–278. https://doi.org/10.1126/science.1229931

Pocock, M.J., Roy, H.E., August, T., Kuria, A., Barasa, F., Bett, J., Githiru, M., Kairo, J., Kimani, J., Kinuthia, W., Kissui, B., Madindou, I., Mbogo, K., Mirembe, J., Mugo, P. Muniale, F.M., Njoroge, P., Njuguna, E.G., Izava, Olendo, M.I., Opige, M., Otieno, T.O., Ng’weno, C.C., Pallangyo, E., Thenya, T., Wanjiru, A., & Trevelyan, R. (2019). Developing the global potential of citizen science: Assessing opportunities that benefit people, society and the environment in East Africa. Journal of applied ecology, 56(2), 274–281. https://doi.org/10.1111/1365-2664.13279

Vanderhoeven, S., Adriaens, T., Desmet, P., Strubbe, D., Backeljau, T., Barbier, Y., Brosens, D., Cigar, J., Coupremanne, M., De Troch, R., Eggermont, H., Heughebaert, A., Hostens, K., Huybrechts, P., Jacquemart, A., Lens, L., Monty, A., Paquet, J., Prévot, C., Robertson, T., Termonia, P., Van De Kerchove, R., Van Hoey, G., Van Schaeybroeck, B., Vercayie, D., Verleye, T., Welby, S., & Groom, Q. (2017). Tracking Invasive Alien Species (TrIAS): Building a data-driven framework to inform policy. Research Ideas and Outcomes, 3: e13414. https://doi.org/10.3897/rio.3.e13414

